# Molecular basis of ligand recognition and activation of human V2 vasopressin receptor

**DOI:** 10.1101/2021.01.18.427077

**Authors:** Fulai Zhou, Chenyu Ye, Xiaomin Ma, Wanchao Yin, Qingtong Zhou, Xinheng He, Xiaokang Zhang, Tristan I. Croll, Dehua Yang, Peiyi Wang, H. Eric Xu, Ming-Wei Wang, Yi Jiang

## Abstract

The V2 vasopressin receptor (V2R) is a class A G protein-coupled receptor (GPCR) and plays a vital role in controlling water homeostasis upon stimulation by the natural peptide arginine vasopressin (AVP). Thus, V2R has attracted intense interest as a drug target for diabetes insipidus, nocturia, and hyponatremia. However, how AVP recognizes and activates V2R remains elusive. Here, we report the 2.6 Å resolution structure of V2R bound to AVP and the stimulatory G protein G_s_, determined by cryo-electron microscopy (cryo-EM). In this complex, AVP presents a unique cyclic conformation formed by an intramolecular disulfide bond and engages the orthosteric binding pocket of V2R in a ligand-specific mode. Comparison of the AVP–V2R–G_s_ complex to previously reported G_s_-coupled class A GPCRs reveals distinct structural features, including a smaller outward movement of TM5 and TM6 and the concomitant shift of G_s_ protein. Our detailed structural analysis provides a framework for understanding AVP recognition and V2R activation, thereby offering a structural template for drug design targeting V2R.

## Introduction

Vasopressin type 2 receptor (V2R) belongs to the vasopressin (VP)/oxytocin (OT) receptor subfamily of G protein-coupled receptors (GPCRs), which comprise at least four closely related receptor subtypes: V1aR, V 1bR, V2R, and OTR^1^. These receptors are activated by arginine vasopressin (AVP, CYFQNCPRG-NH_2_) and OT (CYIQNCPLG-NH_2_), two endogenous nine-amino acid neurohypophysial hormones synthesized in the hypothalamus and secreted from the posterior pituitary gland. AVP and OT are thought to mediate a global biologically conserved role in social behavior and sexual reproduction^2^. Among VP/OT receptors, OT exhibits high selectivity for OTR. Conversely, AVP shows similar affinities to all the four receptors^3^. Upon stimulation by AVP, V2R plays a central role in controlling water homeostasis^4^.

V2R is mainly expressed in the renal collecting duct principal cells and mediates the antidiuretic action of AVP by accelerating water reabsorption. In response to hypovolemia or high osmolality, AVP is secreted from the posterior pituitary and binds to V2R in the basolateral membrane of these principal cells, thus stimulates adenylyl cyclase via the stimulatory G protein (G_s_). This process promotes cAMP/PKA-mediated trafficking of aquaporin-2 (AQP2) water channels to the apical membrane in a long-term manner, allowing water to freely enter via the medullary osmotic gradient^4^. Additionally, ectopic expression of AVP and V2R has been reported in breast, pancreatic, colorectal, and gastrointestinal cancers, as well as renal cell and small cell lung carcinomas, indicated of their roles of tumorgenicitys^5–7^.

Considering its primary role in pathogenesis, V2R is referred to as an attractive drug target for prevention and control of diabetes insipidus, nocturia, hyponatremia secondary to syndrome of inappropriate diuretic hormone secretion (SIADH), and autosomal dominant polycystic kidney disease (ADPKD). Although there has been substantial progress in discovering peptidic and small molecular ligands targeting V2R, desmopressin (dDAVP), a peptidic AVP analogue^8^, is the only agonist approved for the treatment of diabetes insipidus. Additionally, small molecular V2R antagonists, tolvaptan, conivaptan, and atosiban, are presently applied to clinical management of nephrogenic syndrome of inappropriate diuresis (NSIAD) (https://www.cortellis.com/drugdiscovery).

Extensive efforts have been devoted to understanding the mechanism of peptide recognition and activation of V2R by its natural peptide ligands and to develop therapeutic agents via mutagenesis and structure-activity relationship studies^9^ and molecular dynamics simulation^10, 11^. However, except for the structure of OTR bound to a selective small molecular antagonist Retosiban (PDB code: 6TPK)^12^, no other structures from the VP/OT receptor subfamily, especially an agonist-bound active structure, are available. In this study, we determined a near-atomic resolution cryo-EM structure of the full-length, G_s_-coupled human V2R bound to AVP. It reveals a specific recognition mode of AVP by V2R and an unconventional receptor activation mechanism different from that of previously reported G_s_-coupled class A GPCRs, thus providing a structural template for mechanistic understanding and V2R-targeted drug design.

## Results

### Cryo-EM structure of G_s_-coupled V2R bound to AVP

The structure of the AVP-V2R-G_s_ complex was determined to a resolution of 2.6 Å (Fig. 1 and Supplementary information, Table S1). An engineered mini-Gαs protein was used to obtain this complex structure^13^. The N-terminus of the engineered mini-Gα_s_ was replaced by the corresponding residues of Gα_i_ to facilitate the binding of scFv16. An analogous approach was used to obtain the structures of the 5-HT_2A_ receptor bound to Gα_q_ and the Gα_11_-coupled muscarinic M1 receptor^14, 15^.

**Figure 1.**
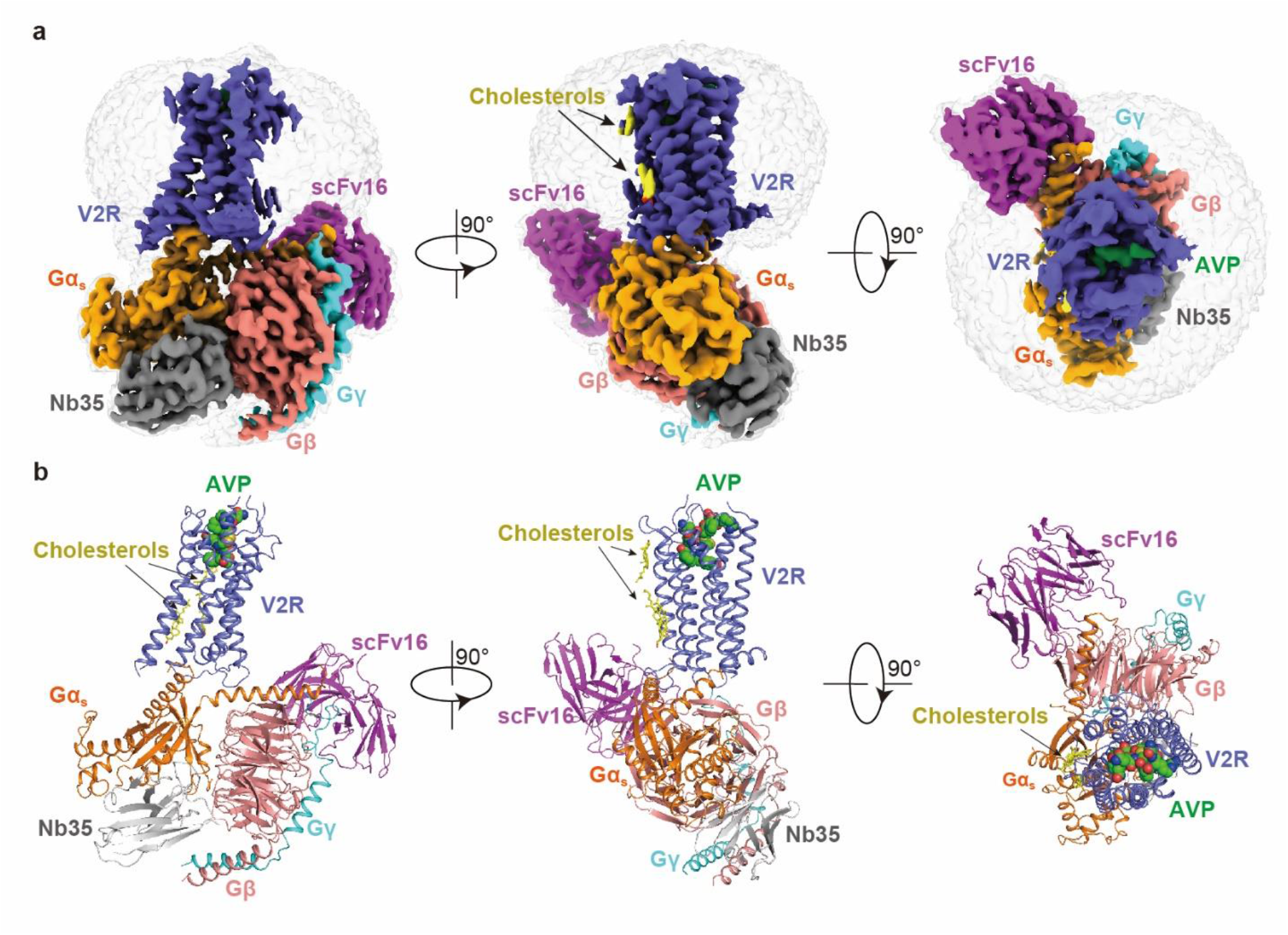
Cryo-EM structure of the AVP–V2R–G_s_ complex. Orthogonal and extracellular views of the density map (**a**) and model (**b**) for the AVP–V2R–G_s_ complex. V2R, slate blue; AVP, arginine vasopressin, green; G_s_, orange; Gβ, salmon; Gγ, cyan; Nb35, gray; scFv16, magenta; cholesterols, yellow.

The final structure of the AVP–V2R–G_s_ complex contains all residues of AVP (residues 1-9), the Gα_s_ Ras-like domain, Gβγ subunits, Nb35, scFv16, and the V2R residues from T31 to L339 ^8.57^ (superscripts refer to Ballesteros–Weinstein numbering^16^) (Fig.1, Supplementary information, Fig. S1). We also used the NanoBiT tethering strategy, which was initially introduced in the structural studies of the class B GPCR-G_s_ complexes^17, 18^, to stabilize the AVP–V2R–G_s_ complex. These modifications have little effect on the pharmacological properties of V2R (Supplementary information, Table S2). The majority of amino acid side chains, including AVP, transmembrane domain (TMD), all flexible intracellular loops (ICLs) and extracellular loops (ECLs) except for ICL3 and G185-G188 in ECL2, were well resolved in the model, refined against the EM density map (Supplementary information, Figs. S2 and S3). Thus, the complex structure can provide detailed information on the binding interface between AVP and helix bundle of the receptor, as well as the interface between G_s_ heterotrimer and the receptor. The TMD is surrounded by an annular detergent micelle mimicking the natural phospholipid bilayer (Fig.1a). Three cholesterol molecules surrounding the GPCR transmembrane domain (TMD) are shown in the final structure (Fig. 1b).

### Molecular recognition of AVP by V2R

AVP occupies an orthosteric binding pocket in the TMD bundle composed of all TM helices and all ECLs (Fig. 2). The most notable conformational feature of AVP is the tocin ring formed by a disulfide bridge between the first and sixth cysteine residues, presenting a “spoon-like” conformation (Fig. 2b). This distinctive architecture is also shared with OT and the synthesized peptide dDAVP^19, 20^, and is the minimal common requirement for the biological activity of these peptides^21^. The cyclic spoon head inserts deeply into the TMD core, while the C- erminus spoon tail stretches toward the ECLs of the receptor (Figs. 2a and b).

**Figure 2.**
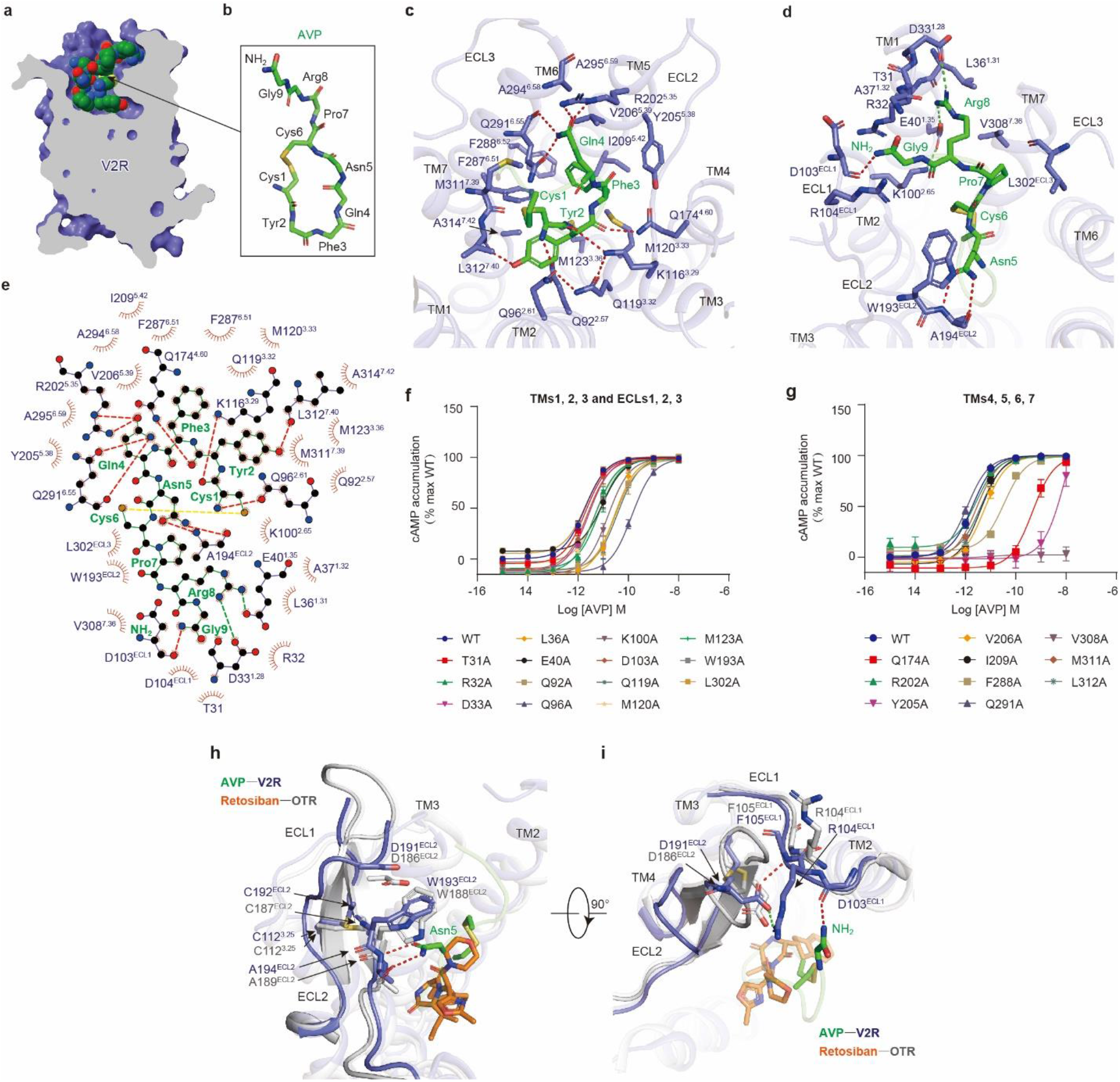
The ligand-binding pocket of V2R. **a**, Vertical cross-section of the AVP binding pocket in V2R. The peptide AVP is shown as spheres with carbons in green. **b**, Spoon-like conformation of AVP. The peptide backbone is shown in green. **c** and **d**, Detailed interactions between V2R and AVP residues (Cys^1P^, Tyr^2P^, Phe^3P^ and Gln^4P^) in (**c**) or AVP residues (Asn^5P^, Cys^6P^, Pro^7P^, Arg^8P^, and Gly^9P^-NH2) in (**d**). H-bonds and salt bridges are depicted as red and green dashed lines, respectively. **e**, Schematic representation of interactions between V2R and AVP analyzed by LigPlot^+^ program^43^. H-bonds and salt bridges are shown as red and green dashed lines, respectively. The disulfide bond between the first and six cysteine residues is shown as a yellow dashed line. The stick drawings and residues labels of V2R and AVP are colored slate blue and green, respectively. **f** and **g**, Effects of mutations in the ligand-binding pocket on AVP-induced cAMP accumulation. cAMP levels were measured in wild type (WT) receptor and alanine mutants in all ECLs, TMs 1, 2 and 3 (**f**), TMs 4, 5, 6 and 7(**g**). Data are shown as means ± S.E.M. of at least three independent experiments conducted in triplicate. **h** and **i**, A comparison of conformational change of ECL1 and ECL2 in AVP-bound V2R and retosiban-bound OTR (PDB code: 6TPK) structures. The receptor and ligand in AVP–V2R structure region are shown in slate blue cartoon and green sticks representation, respectively. The receptor and ligand in OTR structure are shown in gray cartoon and orange sticks representation, respectively. The conformational change of the “DCWA” conserved motif of ECL2 and detailed interactions between “DCWA” motif and AVP are shown (**h**). The polar interaction network involved in D191^ECL2^, F105^ECL1^, and R104^ECL1^ stabilizes ECL1 and ECL2 conformation, thereby facilitating the interactions between ECL1 and the C-terminal amide of AVP (**i**). H-bonds and salt bridge are indicated as red dashed and green lines, respectively. The residues of “DCWA” motif and C112^3.25^ residues are shown as sticks and labeled in AVP–V2R (slate blue) and retosiban–OTR (gray). C187^ECL2^ (OTR) or C192^ECL2^ (V2R) forms a conserved disulfide bond with C112^3.25^, respectively. The disulfide bonds are shown as yellow sticks.

The distinguished cyclic conformation of AVP and the binding pocket’s physicochemical environment render a peptide-specific AVP binding mode (Fig. 2 and Supplementary information, Table S3 and S4). Q96^2.61^ forms a stabilizing H-bond network with the N-terminal amine of Cys^1P^ and the sidechain amine of Q119^3.32^ which in turn stacks against the Tyr^2P^ sidechain. Q119^3.32^ H-bonds with K116^3.29^, which forms an additional H-bond with the backbone CO group of Tyr^2P^. These amino acids (Q96^2.61^, K116^3.29^, Q119^3.32^, and Cys^1P^) form a stabilizing H-bond network (Fig. 2c). The alanine mutations of Q96^1.61^ and Q119^3.32^ significantly diminish the AVP-induced cAMP accumulation, supporting our hypothesis that this H-bond network is critical to AVP-induced V2R activation (Fig. 2f and Supplementary information, Table S3). Tyr^2P^ makes hydrophobic interactions with Q92^2.57^, Q96^2.61^, Q119^3.32^, F287^6.51^, M311^7.39^, L312^7.40^, and A314^7.42^ (Figs. 2c and e). Besides hydrophobic interactions, the hydroxyl group of Tyr^2P^ H-bonds to the main-chain oxygen of L312^7.40^, whereas the main-chain CO group of Tyr^2P^ forms a H-bond with Q174^4.60^ (Figs. 2c and e). Q174^4.60^, together with Q92^2.57^, Q96^2.61^, and Q119^3.32^, make substantial contributions to AVP-induced activation of V2R supported by our alanine mutagenesis analysis (Figs. 2f, g, and Supplementary information, Table S3). Phe^3P^ buries in a hydrophobic cleft constituted by M120^3.33^, M123 ^3.36^, Y205^5.38^, V206^5.39^, I209^5.42^, F287^6.51^, F2 8 8^6.52^, and Q29 1 ^6.55^, of which F288^6.52^ is closely involved in AVP-induced V2R activation (Figs. 2c, e, g, and Supplementary information, Table S3). Other polar contacts are observed between Gln^4P^, R202^5.35^, and the backbone and sidechain oxygens of Q291^6.55^, as well as Asn^5P^ and the main-chain of A194^ECL2^ (Figs. 2d and e). Arg^8P^ forms a part of a complex salt bridge with D33^1.28^, E40^1.32^, and K100^1.32^ (Figs. 2d and e). The K100^1.32^A mutation significantly decreased the AVP-induced cAMP accumulation, although its sidechain carbon atoms make week hydrophobic contacts with the Cys^1P^-Cys^6P^ disulfide bond (Figs. 2f, and Supplementary information, Table S3). Additionally, the C-terminal amide of AVP points to ECL2, stacking against the R104^ECL1^ guanidinium group and H-bonding to the backbone CO group of D103^ECL1^ (Figs. 2d and e).

A “DCWA” sequence in ECL2 is highly conserved throughout VP/OT receptor sub-family but is not a feature of other class A GPCRs^22^. In this conserved sequence, W193^ECL2^ and A194^ECL2^ directly contact with AVP, of which W193^ECL2^ anchors Asn^5P^ as a lid (Fig. 2h). Interestingly, when comparing structures of the AVP– V2R-G_s_ complex with that of OTR in inactive state^12^, a steric clash can be observed between Asn^5P^ and W188^ECL2^ of OTR, the cognate residue of W193^ECL2^ in V2R, which may probably confer a notable conformational transition of ECL2 from β-hairpin to a flexible loop upon AVP binding (Fig. 2h). Although D191^ECL2^ does not directly interact with AVP, it may form a H-bond to the F105^ECL1^ amide and a salt-bridge to R104^ECL1^, which in turn, stabilize ECL1 and ECL2 conformation, thereby facilitating the interactions between ECL1 and the C-terminal amide of AVP (Fig. 2i). The significance of D191^ECL2^ to AVP-induced V2R activation is supported by our alanine mutagenesis analysis (Supplementary Table S3). The disulfide bond conserved across class A GPCRs is formed between C194^ECL2^ located in this sequence and C112^3.25^ and may contribute to the stabilization of ECL2 conformation (Fig. 2i).

This structure also provides a template to study the selectivity of AVP and OT (Supplementary information, Fig. S4). OT has two amino acid substitutions at positions 3 and 8, Phe^3P^ to Ile^3P^, and Arg^8P^ to Leu^8P^, respectively (Supplementary information, Fig. S4a). It exhibits a much lower selectivity against V2R, with an over 500-fold decrease in binding affinity as opposed to AVP^3^. Interestingly, residues surrounding Phe^3P^ and Arg^8P^ are closely associated with AVP-induced V2R activation (Supplementary information, Table. S3), indicating the environment faced by amino acids at positions 3 and 8 are probably critical to OTR activation. By docking OT into the V2R binding pocket, a weaker hydrophobic interaction network is created for Ile^3P^ compared to its equivalent residue Phe^3P^ in AVP (Supplementary information, Fig. S4b). Additionally, the hydrophobic leucine at position 8 breaks the polar interactions between Arg^8P^ of AVP and receptor (Supplementary information, Fig. S4c). Besides, Leu^8P^ faces a relatively weak hydrophobic environment composes of R32, D33^1.28^, L36^1.31^, A37^1.32^, and E40^1.35^. Thus, the peptide-binding pocket of V2R defines a relatively unfavorable binding environment for OT in contrast to AVP, which may hamper OT binding.

Collectively, AVP interacts with V2R in a ligand-specific manner. These observations provide a rationale for understanding V2R recognition by AVP and OT, as well as a structural template for V2R-targeted drug discovery.

### Activation of V2R by AVP

A comparison of the AVP–V2R–G_s_ complex structure to that of OTR in the inactive state sheds light on the conformational changes involved in activation of V2R: The cytoplasmic end of TM6 in G_s_-coupled V2R undergoes a notable outward displacement, the hallmark of GPCR activation (Fig. 3a). V2R conforms to a common activation pathway that directly links the ligand-binding pocket to G-protein coupling regions in class A GPCRs^23^, including the AVP-induced turning of the rotamer “toggle switch” W284^6.48^, which translates into the rotation and outward movement of TM6 (Fig. 3b). Several distinct features on active V2R conformation can be observed compared to other previously reported class A GPCR structures, suggesting that V2R may undergo a receptor-specific activation.

**Figure 3.**
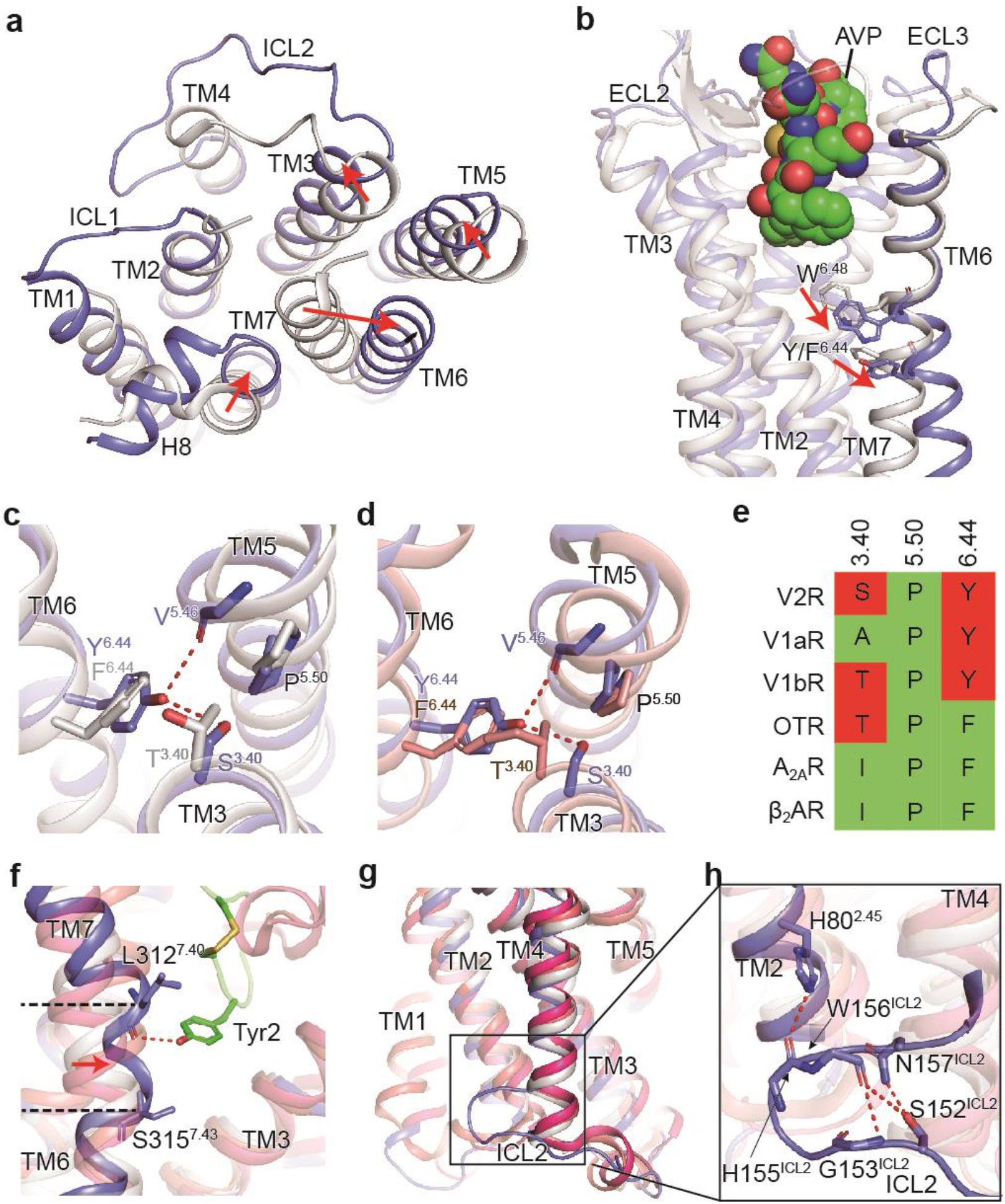
Activation mechanism of V2R. **a**, The cytoplasmic view of the transmembrane helical bundles in AVP-bound V2R (slate blue) and retosiban-bound OTR (gray) structures shown in cartoon representation. The red arrows indicate the movements of transmembrane helices (TM3, TM5, TM6, and TM7) in the active V2R structure relative to the inactive OTR structure. **b**, Conformational changes of W^6.48^ and Y/F^6.44^. The structures of AVP–V2R (slate blue) and retosiban–OTR (gray) are shown as cartoons, and AVP is shown as green spheres. The red arrows indicate the movement of the side chain of W^6.48^ and Y/F^6.44^ (shown as sticks) in the AVP–V2R (active) structure compared to retosiban–OTR (inactive) structure. **c** and **d**, Conformational changes of the conserved P^5.50^I^3.40^F^6.44^ motif in AVP-bound V2R structure relative to inactive OTR (**c**) and active β_2A_R structures (**d**), respectively. The structures are shown as cartoon and colored in slate blue (V2R), gray (OTR), and deep salmon (β_2A_R). Sidechains of P^5.50^, S^3.40^, Y^6.44^, and main-chain of V^5.46^ in V2R, side chains of P^5.50^, T^3.40^, F^6.44^ in OTR and P^5.50^, I^3.40^, F^6.44^ in β_2A_R are shown as sticks. The stick drawings and residues labels of V2R, OTR and β_2A_R are colored in slate blue, gray and green, respectively. H-bonds are depicted as red dashed lines. **e**, Sequence comparison of conserved p^5.50^-I^3.40^-F^6.44^ motif between V2R, V1aR, V1bR, OTR A_2A_R, and β_2A_R. Polar residues are colored in red and nonpolar residues are colored in green. **f**, Conformational change of TM7 in AVP–V2R structure relative to retosiban–OTR (gray), β_2A_R–G_s_ (deep salmon), and A_2A_R–G_s_ (hot pink) structures shown as cartoons. AVP is shown as green cartoon with its Tyr^2P^ side chain displayed as sticks and intramolecular disulfide bond as yellow stick. The distortion range was indicated by two dark dashed lines pointed to the residues L312^7.40^ and S315^7.43^ shown as sticks. Red arrow indicates the movement of TM7 in V2R. The H-bond is depicted as a red dashed line. **g**, Conformational change of the cytoplasmic end of TM4 in AVP–V2R structure relative to retosiban–OTR (gray), β_2A_R–G_s_ (deep salmon), and A_2A_R–G_s_ (hot pink) structures shown as cartoons. **h**, Close view of the C-shaped conformational change of TM4 in AVP–V2R structure. H80^2.45^, H155^ICL2^, and the main-chain of G153^ICL2^, H155^ICL2^, W156^ICL2^ consisting of a polar interaction network in the C-shaped loop are shown as sticks. H-bonds and polar interactions are depicted as red dashed lines, respectively.

To the best of our knowledge, residues P^5.50^, I^3.40^, and F^6.44^, the conserved “P-I-F” micro-switch, undergoes a conformation rearrangement during receptor activation^24^. Indeed, this “P-I-F” micro-switch of V2R exhibits a substantial rearrangement compared to inactive OTR, reflecting the activation state of V2R (Fig. 3c). The sequence alignment analysis shows that the residue F^6.44^ is evolutionally conserved among 74% of class A GPCRs, while the hydrophobic residues Ile/Leu/Val account for residues at position 3.40 in 74% of class A GPCRs^25^, thus providing a hydrophobic environment for the packing of TM3-5-6. By contrast, F’^3.44^ and I^3.40^ are substituted by polar residues Y280^6.44^ and S127^3.40^ in V2R, respectively (Figs. 3d and e). Compared to the G_s_-coupled β_2_ adrenergic receptor (β_2A_R, PDB code: 3SN6)^26^, the distinct physicochemical environment in V2R facilitates the formation of H-bonds between Y280^6.44^ and S127^3.40^, as well as Y280^6.44^ and the backbone CO group of V213^5.46^, which probably stabilize the active conformation of the receptor (Fig. 3d). The disease-associated mutations S127^3.40^F and Y280^6.44^C deactivate V2R, offering experimental evidence to support these residues’ putative role in V2R activation^27, 28^. The alanine mutation of F284^6.44^ abolished the binding and activation of OTR, while the replacement of F284^6.44^ in OTR by the V2R-equivalent tyrosine slightly decreased the V2R activation by AVP but converted AVP from a partial to a full agonist^29^. These data are consistent with the hypothesis that polar interactions among these unconventional “P-I-F” residues may differentiate the activation mode between V2R and OTR.

Intriguingly, a distortion of TM7 helix was observed between L312^7.40^ and S315^7.43^, demonstrating a unique structural feature that is different from other class A GPCRs solved so far (Fig. 3f). This conformation appears to be stabilized by the H-bond between Tyr^2P^ of AVP and the main-chain CO group of L312^7.40^ in the receptor. This polar contact draws TM7 closer to the core of the peptide-binding pocket (Fig. 3f). [Phe2]AVP, a synthesized AVP analog with a substitution of Tyr by Phe abolishing this H-bond, decreased V2R activity by 28-fold^30^, implying that the polar interaction between Tyr^2P^ and the receptor is critical to AVP-induced V2R activation.

Furthermore, the cytoplasmic end of TM4 is one helical turn shorter than other G_s_-coupled GPCRs, thereby releasing a longer ICL2 (Fig. 3g). The latter protrudes towards TM1 and adopts a C-shaped conformation (Fig. 3g). This distinct conformation is stabilized by a H-bond formed between H80^2.45^ and the backbone CO group of H155^ICL2^ (Fig. 3h). The C-shaped loop is further stabilized by a polar interaction network constituted by the side chains of N157^ICL2^ and S152^ICL2^, as well as the backbones of G153^ICL2^ and W156^ICL2^ (Fig. 3h). These interactions may stabilize ICL2 in a specific conformation and probably affect the ICL2-Gα_s_ interaction pattern.

### G protein coupling by V2R

The interface between V2R and G_s_ heterotrimer consists of four transmembrane helices (TM2, TM3, TM5, and TM6), ICL2, and helix 8. The outward movement of the cytoplasmic ends of TM5 and TM6 open a cytoplasmic cavity together with TM2, TM3, and helix 8 to accommodate the α5 helix of Gα_s_. Compared to other solved G_s_-coupled class A GPCRs, the AVP-V2R-G_s_ complex shows a distinctive GPCR-G_s_ coupling feature (Fig. 4a and b). The cytoplasmic ends of TM5 and TM6 in V2R display a smaller amplitude of outward displacements than the G_s_-coupled β_2A_R (PDB code: 3SN6; 5.8 Å for TM5 and 8.7 Å for TM6)^26^ and adenosine A_2A_ receptor (A_2A_R, PDB code: 5G53; 2.5 Å for TM5 and 7.2 Å for TM6 measured at Cα carbon of residues 5.67 and 6.30, respectively) (Fig. 4a)^31^. In addition, the α5 helix of Gα_s_ in the AVP–V2R–G_s_ complex shifts 0.5 helical turns away from the 7TM core (Fig. 4a). These structural differences altogether create a smaller cytoplasmic cavity to accommodate G_s_ protein. Indeed, the solvent-accessible surface area (SASA) of the V2R-Gα_s_ interface (943.32 Å^2^) is smaller than that of β_2A_R-Gα_s_ (1,030 Å^2^) and A_2A_R-G%_s_ interfaces (1,276 Å^2^) (Fig. 4c). A noticeable shift of G_s_ protein in the AVP–V2R–G_s_ complex occurs (Fig. 4a and b) that may partly be caused by relatively inward positions of TM5 and TM6, thus pushing the entire G_s_ heterotrimer shifts in the same direction.

**Figure 4.**
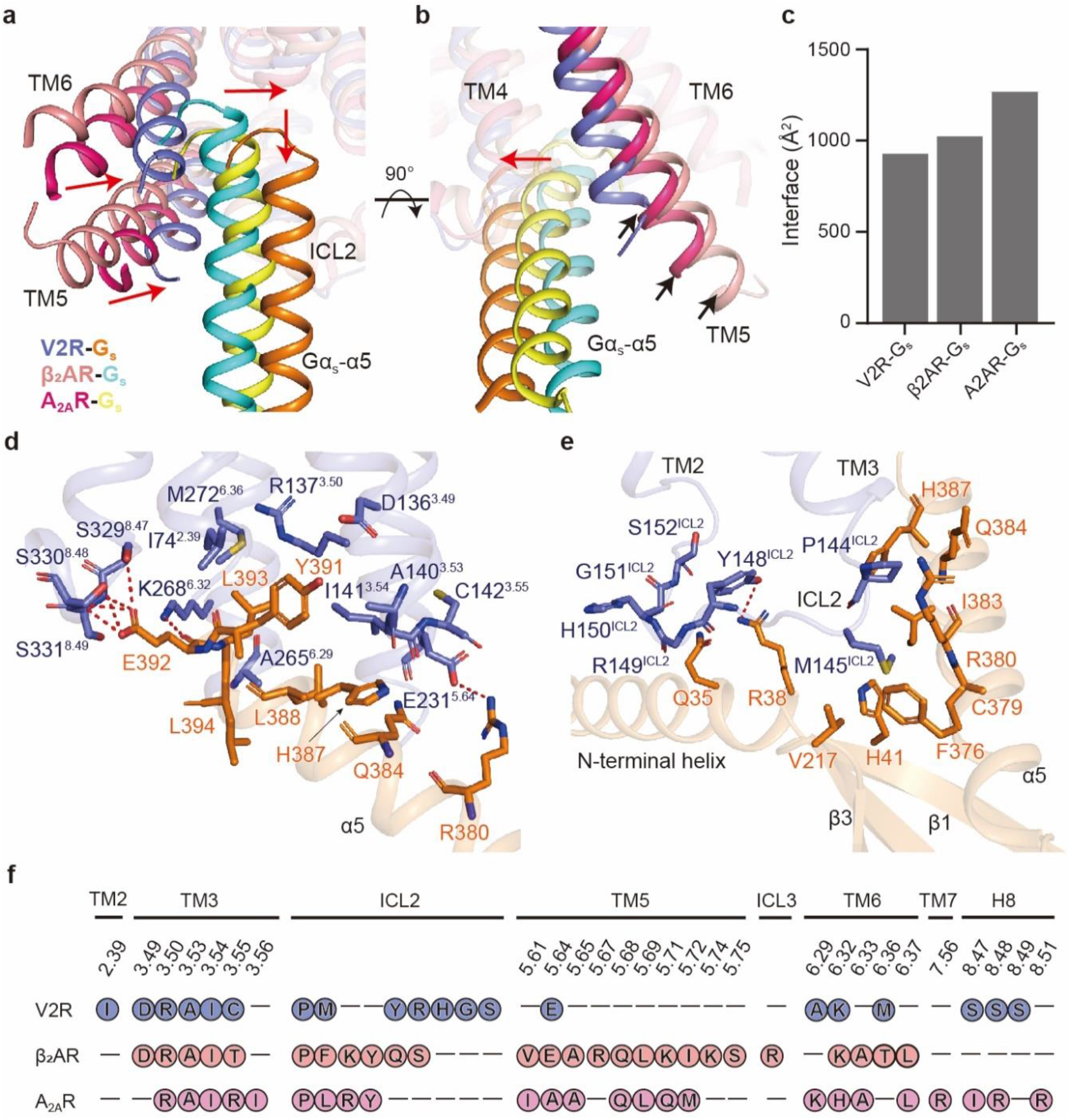
G_s_ protein-coupling of V2R. **a**, A comparison of receptor helical bundle and Gα_s_ α5 helix conformations between AVP–V2R–G_s_ and two other class A GPCR–G_s_ complexes. AVP–V2R–G_s_, β_2A_R–G_s_, and A_2A_R–G_s_ are colored in slate blue, deep salmon, and hot pink for the receptor, and orange, cyan, and yellow for G_s_ protein, respectively. Red arrows indicate the movements of intracellular tips of helices 5, 6 and α5 helix of Gα_s_ from V2R–G_s_ complex compared to β_2A_R–G_s_ or A_2A_R–G_s_ complexes. **b**, A comparison of TM5 length and shift of α5 helix of Gα_s_. Red arrow indicates the shift of Gα_s_ α5 helix. Black arrows show the cytoplasmic ends of TM5 in the AVP–V2R–G_s_, β_2A_R–G_s_, and A_2A_R–G_s_ structures. **c**, Calculated solvent-accessible surface area (SASA) of the interfaces between receptors and G_s_ protein in Chimera X^44^. **d** and **e**, Detailed interactions of helices (TM2, TM3, TM5, TM6, and H8) with α5 helix of G_s_ (**d**) and ICL2 with N-terminal and α5 helices, β1 and β3 strands of G_s_ (**e**). H-bonds are depicted as red dashed lines. The stick drawings and residues labels of V2R and Gα_s_ are colored in slate blue and orange, respectively. **f**, Residues in V2R (slate blue), β_2A_R (deep salmon), and A_2A_R (hot pink) that contact with G_s_.

The major binding interface between V2R and G_s_ protein is between the α5 helix of the Gα_s_ subunit and the cytosolic core formed by TMD (Fig. 4d). In this interface, E392 of Gα_s_ polar interacts with three consecutive serines in receptor helix 8 (S3 2 9^8.47^, S330^8.48^, and S331^8.49^). Another polar interaction can be observed between R380 and E231^5.64^. These polar interactions may contribute to stabilizing the V2R-Gα_s_ interface, which is crucial to GPCR-G protein coupling. One additional interface relates to ICL2 that interacts with α5 helix, αN-β1 junction, and the top of the β3-strand of Gα_s_, presumably stabilized by hydrophobic contacts (Fig. 4e).

The GPCR-G_s_ coupling profiles of V2R, β_2A_R, and A_2A_R point to several distinct features (Fig. 4f). ICL2 of V2R makes more extensive hydrophobic interactions with Gα_s_ compared to β_2A_R and A_2A_R (Fig. 4f). The superposition of the three also exhibits a 1-2 helical turn shorter TM5 cytoplasmic end in V2R (Fig. 4b). Since proline and glycine are well-known α-helix breakers that disrupt α-helical backbone conformation’s regularity, a consecutive P-G-P sequence (P238, G239, and P240) may terminate the extension of the TM5 (Supplementary information, Fig. S5). Thus, compared to G_s_-coupled β_2A_R and A_2A_R, a shorter TM5 in V2R produces fewer interactions with Gα_s_ (Fig. 4b and f).

### Implication of disease-causing mutations

Numerous mutations of V2R were identified and closely associated with human diseases^32^. Gain-of-function mutations in the gene encoding the V2R *(AVPR2)*, including R137^3.50^C/L, F229^5.62^V, and I130^3.43^N, cause NSIAD, leading to hyponatremia and related clinical symptoms in infants^33–35^. Conversely, loss-of-function mutations of *AVPR2* account for almost 90% incidence rate of X-linked congenital nephrogenic diabetes insipidus (NDI)^36, 37^.

These reported disease-associated mutations are located in three major regions of V2R: the ligand-binding pocket, the G protein-coupling site, and the central region connecting these two regions (Supplementary information, Fig. S6)^23, 32^. The activating L312^7.40^S mutation and inactivating mutations R202^5.35^C, F287^6.51^L, A294^6.58^P, and M311^7.39^V all sit in the AVP binding pocket. It is notable that L312^7.40^ sits at the site of the unique distortion of TM7 mentioned above. Substitutions of residues in ECLs to cysteines (R104^ECL1^C, R106^ECL1^C, and R181^ECL2^C) led to loss-of-function mutations^38^, which are putatively attributed to incorrect disulfide bonds formation by these cysteines^39^. A majority of disease-associated mutations occur in the central region linking the bottom of the ligand-binding pocket with the G protein-coupling site. Several mutations arise at the highly conserved micro-switches (S127^3.40^F and Y280^6.44^C in P-I-F, N321^7.49^K/Y and Y325^7.53^A in NPxxY, R137^3.50^H/C/L in D/ERY, as well as D85^2.50^N and N317^7.45^S in Na^+^ pocket). Not surprisingly, these mutations can affect V2R activation since they are related to key residues responsible for subsequent G protein-coupling^23^. In the G protein-coupling region, R137^3.50^ and S329^8.47^ directly interact with G_s_ protein in our structure. Interestingly, mutations in R137^3.50^ can result in opposite phenotypes (R137^3.50^C/L for gain-of-function and R137^3.50^H for loss-of-function), probably due to the mutation-elicited changes in the physicochemical environmentunsuitable to TM association and G_s_ coupling.

Collectively, our findings provide a structural basis for V2R mutation-associated diseases. It is worth noting that the mechanisms involved are complex. Besides functional disorder, mislocalization by impaired intracellular trafficking and abnormal receptor expression disorder are also implicated in diseases. Finally, most V2R mutations are likely associated with combined disorders^32^. Systematic investigations are required to enrich our knowledge in this regard.

## Discussion

The near-atomic resolution single-particle cryo-EM structure of G_s_-coupled V2R bound to its endogenous ligand AVP reported in this study shows that the peptide adopts a “spoon-like” conformation with its featured tocin ring inserting into the TMD bundle and the C-terminus facing the binding pocket. This tocin ring is connected by a disulfide bridge formed between the first and sixth cysteine residues. Obviously, the cyclic backbone is shared with OT. It is conceivable that the distinct physicochemical environment in the peptide-binding pocket may lead to the low selectivity of OT to V2R. Our findings thus provide a framework for a better understanding of ligand recognition by V2R. Structural comparison with other G_s_-coupled class A GPCRs sheds light on the basis of AVP-induced V2R activation. Although V2R conforms to a common activation mechanism, AVP-induced distortion of TM7 and a unique polar network formed by equivalent residues in the “P-I-F” motif may differentiate activation of V2R and OTR from other class A GPCRs. A smaller amplitude of the outward displacement of TM5 and TM6, as well as the concomitant shift of Gα_s_, not only distinguish V2R from its counterparts but also represent a diversified G_s_ coupling mechanism. Of note is that ICL3 of V2R/OTR was shown to be a determinant in G protein coupling^40^. Considering ICL3 is invisible in our structure, this conclusion deserves verification by additional structural information.

Our findings also offer a structural template for designing better ligands targeting V2R. dDAVP is an AVP analog with moderate selectivity to V1b and OT but is devoid of V1a binding. It has a sound antidiuretic effect. However, it also increases the risk of hyponatremia caused by a long plasma half-life and the resulting clearance delay from the kidney^41^. Efforts have been made to design potent V2R agonists with improved selectivity and pharmacokinetic profiles^42^. Such an endeavor will certainly be benefited from the structure information present in this paper.

## Materials and Methods

### Cell culture

*Spodoptera frugiperda* 9 (*Sf*9) insect cells (Expression Systems) were grown in ESF 921 serum-free medium (Expression Systems) at 27°C and 120 rpm. HEK 293T cells were purchased from American Type Culture Collection (ATCC), cultured in Dulbecco’s modified Eagle’s medium (DMEM, Life Technologies) supplemented with 10% fetal bovine serum (FBS, Gibco), and maintained in a humidified chamber with 5% CO2 at 37°C.

### Constructs of V2R and G_s_ heterotrimer

The human wild-type (WT) V2R gene was codon-optimized for *Sf9* expression and synthesized by Sangon Biotech. To facilitate the expression and purification, the full-length V2R DNA was cloned into a modified pFastBac vector (Invitrogen), which contains an N-terminal hemagglutinin (HA) signal peptide followed by a 10 × His-tag and a b562RIL (BRIL) epitope before the receptor. To improve the homogeneity and stability, the NanoBiT tethering strategy was applied by fusing a LgBiT subunit (Promega) at the receptor C-terminus using homologous recombination (CloneExpress One Step Cloning Kit, Vazyme) (Supplementary information, Fig. S1)^17^.

An engineered G_s_ construct (G112) was designed based on mini-G_s_ that was used in the crystal structure determination of A_2A_R–mini-G_s_ complex^31^. By replacing the N-terminal histidine tag (His6) and TEV protease cleavage site with the N-terminal eighteen amino acids (M1-M18) of human G_i1_, this chimeric G_s_ was capable of binding to scFv16, which was used to stabilize the GPCR–G_i_ or –G_11_ complexes^14^–^45^. Additionally, replacing the GGSGGSGG linker at the position of original Gα_s_ α-helical domain (AHD, V65-L203) with that of human G_i1_ (G60-K180) provided the binding site for Fab_G50, another antibody fragment which was used to stabilize the rhodopsin-Gi complex^46^. Furthermore, three mutations (G226A, L272D, and A366S) were incorporated through site-directed mutagenesis as previously described to further increase the dominant-negative effect by stabilizing the Gαβγ heterotrimer^47, 48^. These modifications enabled the application of different nanobodies or antibody fragments to stabilize the receptor-G_s_ complex, although Nb35 and scFv16 were used together during the AVP–V2R–G_s_ complex formation and stabilization in this study. Rat Gβ1 was modified to fuse a SmBiT subunit^17, 49^ (peptide 86 and also named as HiBiT, Promega) with a 15-amino acid (15AA) polypeptide linker (GSSGGGGSGGGGSSG) at its C-terminus. The engineered G_s_ (G112), Gβ1-15AA-HiBiT, and bovine Gγ2 were cloned into the pFastBac vector, respectively. scFv16 was cloned into a modified pFastBac vector which contains a GP67 secretion signal peptide at its N-terminus. For constructs used in functional assays, they were all cloned into the pcDNA3.1 vector (Invitrogen) with an N-terminal Flag (DYKDDDD) tag proceeded by a HA signal sequence.

### Expression and purification of nanobody35

Nanobody35 (Nb35) with a C-terminal 6 × His-tag was expressed in *E.coli* BL21 (DE3) bacteria and cultured in TB medium supplemented with 2 mM MgCl_2_, 0.1% (w/v) glucose and 50 μg/mL ampicillin to an OD_600_ value of 1.0 at 37°C, 180 rpm. Nb35 expression were then induced by adding 1 mM IPTG into the cultures and grown for 5 h at 37°C. Cells were then harvested by centrifugation (4,000 rpm, 20 min) and Nb35 protein was extracted and purified by nickel affinity chromatography as previously described^50^. Eluted protein was concentrated and subjected to a HiLoad 16/600 Superdex 75 column (GE Healthcare) pre-equilibrated with buffer containing 20 mM HEPES, pH 7.5 and 100 mM NaCl. The monomeric fractions supplemented with 30% (v/v) glycerol were flash frozen in liquid nitrogen and stored in −80°C until use.

### Expression and purification of the AVP–V2R–G_s_ complex

Baculoviruses were prepared using the Bac-to-Bac Baculovirus Expression System (Invitrogen). *Sf9* insect cells were cultured to a density of 3 × 10^6^ cells per mL and co-infected with His10-BRIL-V2R-LgBiT, engineered G_s_ (G112), Gβ1-15AA-HiBiT, Gγ2, and scFv16 baculoviruses at a 1:1:1:1:1 ratio. The cells were then harvested by centrifugation 48 h post-infection and stored in −80°C for future use.

The frozen cells were thawed on ice and resuspended in lysis buffer containing 20 mM HEPES, pH 7.5, 100 mM NaCl, 10% (v/v) glycerol, 10 mM MgCl_2_, 5 mM CaCl_2_, 100 μM TCEP (Sigma-Aldrich) and supplemented with EDTA-free protease inhibitor cocktail (Bimake). Cells were lysed by dounce homogenization and complex formation was initiated with the addition of 10 μg/mL Nb35, 25 mU/mL Apyrase (Sigma-Aldrich) and 10 μM vasopressin (GenScript) for 1.5 h at room temperature (RT). The membrane was then solubilized by adding 0.5% (w/v) lauryl maltose neopentyl glycol (LMNG, Anatrace) and 0.1% (w/v) cholesterol hemisuccinate (CHS, Anatrace) for 2 h at 4°C. The sample was clarified by centrifugation at 30,000 rpm for 30 min and the supernatant was then incubated with TALON resin (Clontech) supplemented with 10 mM imidazole for 3 h at 4°C. After incubation, the resin was collected by centrifugation (600 *g*, 10 min) and loaded into a gravity flow column, followed by first wash with five-column volumes of 20 mM HEPES, pH 7.5, 100 mM NaCl, 10% (v/v) glycerol, 5 mM MgCl_2_, 2 μM vasopressin, 25 μM TCEP, 10 mM imidazole, 0.1% (w/v) LMNG and 0.02% (w/v) CHS and then, with fifteen-column volumes of 20 mM HEPES, pH 7.5, 100 mM NaCl, 10% (v/v) glycerol, 5 mM MgCl_2_, 2 μM vasopressin, 25 μM TCEP, 25 mM imidazole, 0.03% (w/v) LMNG, 0.01% (w/v) glyco-diosgenin (GDN, Anatrace), and 0.008% (w/v) CHS. The protein was finally eluted with ten-column volumes of 20 mM HEPES, pH 7.5, 100 mM NaCl, 10% (v/v) glycerol, 5 mM MgCl_2_, 2 μM vasopressin, 25 μM TCEP, 200 mM imidazole, 0.03% (w/v) LMNG, and 0.01% (w/v) GDN. The purified AVP–V2R–G_s_ complex was concentrated using a Amicon Ultra centrifugal filter (molecular weight cut-off of 100 kDa, Millipore) and then subjected to a Superdex 200 Increase 10/300 GL column (GE Healthcare) that was pre-equilibrated with buffer containing 20 mM HEPES, pH 7.5, 100 mM NaCl, 2 mM MgCl_2_, 100 μM TCEP, 5 μM vasopressin, 0.00075% (w/v) LMNG, 0.00025% (w/v) GDN, 0.0002% (w/v) digitonin (Anatrace), and 0.0002% (w/v) CHS. The monomeric fractions of the complex were collected and concentrated to 4-5 mg/mL for cryo-EM examination.

### Cryo-EM data collection and images processing

The freshly purified AVP–V2R–G_s_ complex (3.0 μL) at a final concentration of 4.2 mg/mL was applied to glow-discharged holey carbon grids (Quantifoil R1.2/1.3, 300 mesh), and subsequently vitrified using a Vitrobot Mark IV (ThermoFisher Scientific). Cryo-EM images were collected on a Titan Krios microscope (ThermoFisher Scientific) equipped with a K2 Summit direct electron detector (Gatan) and a GIF Quantum energy filter (slit width 20 eV) in the Cryo-Electron Microscopy Research Center at Southern University of Science and Technology. A total of 4,796 movies were automatically acquired using SerialEM^51^ in super-resolution counting mode at a pixel size of 0.42 Å and with a defocus values ranging from −0.5 to −2.0 μm. Movies with 32 frames each were collected at a dose of 8 electrons per pixel per second over an exposure time of 5.28 s, resulting in an accumulated of dose of 60 electrons per Å^2^ on sample.

All cryo-EM data were processed using RELION-3.1^52^ as shown in the flowchart of Supplementary Fig. S2. Dose-fractionated image stacks were subjected to beam-induced motion correction and dose-weighting using MotionCor2.1^53^. The Contrast transfer function (CTF) parameters for each micrograph were determined by CTFFIND-4.1^54^ and micrographs whose maximum estimated resolution was worse than 4 Å were excluded, leaving 4,611 micrographs for further processing. Particle selection, two-dimensional (2D) and three-dimensional (3D) classifications were performed on a binned dataset with a pixel size of 0.84 Å. Auto-picking yielded 2,583,920 particle projections which were extracted downscaled and subjected to two-rounds of reference-free 2D classification to discard particles with false positives or poorly defined classes, resulting in 1,762,197 particles for 3D processing. This subset of particles was selected and re-extracted without downscaling to produce a 3D initial model for further consecutive rounds of 3D classification. With a 40 Å low-pass filter of initial model, two rounds of 3D classification yielded one well-defined subset of 577,084 particles to be subjected to CTF refinement and three-rounds Bayesian Polishing before the final 3D auto-refinement and sharpening was applied. The final refinement generated a map with an indicated global resolution of 2.6 Å, with 577,084 projections at a Fourier shell correlation of 0.143. Local resolution was determined using the Resmap package with half maps as input maps^55^.

### Model building and refinement

The density map was automatic post-processed using DeepEMhancer^56^ to improve the EM map quality before model building. The initial V2R model was generated by an online homology modeling server, SWISS-MODEL^57^, using the oxytocin receptor (OTR) structure (PDB code: 6TPK) as a template. The mini-G_s_ heterotrimer (mini-G_s_, Gβ1 and Gγ2) and Nb35 taken from GPR52–mini-G_s_–Nb35 complex (PDB code: 6LI3), scFv16 taken from CB2–G_i_–scFv16 complex (PDB code: 6PT0) and vasopressin taken from typsin-vasopressin complex (PDB code: 1YF4) were used as initial models. All models were fitted into the EM density map using UCSF Chimera^58^, followed by iterative rounds of manual adjustment and automated rebuilding in COOT^59^ and PHENIX^60^, respectively. The model was finalized by rebuilding in ISOLDE^61^ followed by refinement in PHENIX with torsion-angle restraints to the input model. The final model statistics were validated using Comprehensive validation (cryo-EM) in PHENIX^60^ and provided in the supplementary information, Table S1. All structural figures were prepared using Chimera^58^, Chimera X^44^ and PyMOL (https://pymol.org/2/).

### cAMP accumulation assay

HEK 293T cells were transiently transfected with different V2R constructs: HA-Flag-V2R(1-371), HA-Flag-His10-BRIL-V2R(1-371)-LgBiT, or HA-Flag-V2R mutants. The cells were digested by 0.02% (w/v) EDTA, resuspended by stimulation buffer (1 × HBSS, 5 mM HEPES, 0.5 mM IBMX, and 0.1% (w/v) BSA, pH 7.4) 24h after transfection, and seeded at a density of 3,000 cells per well into 384-well plates (PerkinElmer). Cells were stimulated by different concentrations of AVP for 40 min at RT. The reaction was terminated by addition of 5 μL Eu-cAMP tracer and 5 μL ULight-anti-cAMP (diluted by cAMP detection buffer). After incubation at RT for 1h, cAMP signals were detected by measuring the fluorescence intensity at 620 nm and 650 nm using an EnVision multilabel plate reader (PerkinElmer).

### Receptor surface expression

Cell-surface expression levels of WT or mutants V2R were quantified by flow cytometry. HEK 293T cells were seeded at a density of 6 ×10^5^ per well into 6-well culture plates. They were digested 24 h post-transfection by 0.02% (w/v) EDTA and blocked with 5% (w/v) BSA in PBS at RT for 15 min, followed by the incubation with 1:300 diluted anti-Flag primary antibody (Sigma) at RT for 1 h. Cells were then washed by 1% (w/v) BSA in PBS three times followed by incubation 1:1000 donkey-anti-mouse Alexa Fluor 488 conjugated secondary antibody (Invitrogen) at 4°C in the dark. After another three washes, cells were resuspended by 200 μL 1% (w/v) BSA in PBS for detection by a NovoCyte flow cytometry (Agilent).

### Molecular docking

The entire process was done in Schrodinger Suite 2017-4. The AVP–V2R–G_s_ complex structure was prepared by Protein Preparation Wizard^62^, hydrogen atoms were added using Maestro and protonation states were assigned using PROPKA. The directions of polar hydrogens were optimized and the whole structure was minimized. Because the peptides to be analyzed have over 100 atoms thus are not suitable for traditional molecular docking, we performed MMGBSA^63^ calculation to estimate relative binding affinity for a list of congeneric ligands using the Prime MM-GBSA module. The analog of AVP, OT was gradually mutated on the basis of AVP after several rounds of energy minimization. Besides, the residues within 4 Å of peptides were further subjected to relax with the sampling method “Minimize” in the Prime MM-GBSA module.

### Statistical analysis

All functional data were analyzed using Prism 7 (Graphpad) and presented as means ± S.E.M. from at least three independent experiments. Concentration–response curves were evaluated with a three-parameter logistic equation to determine the pEC_50_ and *E_max_* values. The significance was determined by one-way ANOVA in Prism 7 and *P* value < 0.01 was considered statistically significant.

## Acknowledgements

We thank all staff members of the Cryo-EM Centre, Southern University of Science and Technology for their assistance in data collection. This work was partially supported by the National Natural Science Foundation of China (31770796 to Y.J., 81872915 to M,-W.W., 31600606 to X.Z., and 81773792 to D.Y.); the National Science & Technology Major Project “Key New Drug Creation and Manufacturing Program” (2018ZX09711002-002-002 to Y.J., 2018ZX09735-001 to M.-W.W., and 2018ZX09711002-002-005 to D.Y.); Ministry of Science and Technology (China) grant (2018YFA0507002 to H.E.X.); Shanghai Municipal Science and Technology Major Project (2019SHZDZX02 to H.E.X.); CAS Strategic Priority Research Program (XDB37030103 to H.E.X.); Start-up funding by Fudan University (Q.Z.); Wellcome Trust (209407/Z/17/Z); the National Key R&D Program of China (2016YFA0501100 to X.Z.); Guangdong Provincial Key Laboratory of Brain Connectome and Behavior (2017B030301017 to X.Z.); CAS Key Laboratory of Brain Connectome and Manipulation (2019DP173024 to X.Z.)

## Author contribution

F.Z. designed the expression constructs, purified the AVP–V2R–G_s_ complex, prepared the final samples for negative stain, cryo-EM grid preparation, and data collection toward the structure, and participated in figure and manuscript preparation; F.Z. and X.M. performed specimen screening by negative-stain EM, cryo-EM data collection, and map calculations; F.Z., X.M. X.Z. built the structure model; F.Z. and T.C. refined the structure model; C.Y. conducted functional studies under the supervision of D.Y. who also participated in data analysis and manuscript editing; Q.Z. and X.H. carried out docking analysis and participated in manuscript preparation; W.Y. engineered the mini-G_s_ protein; F.Z. and Y.J. prepared the bulk of figures, performed the structural analysis, and drafted the manuscript; Y.J., H.E.X., and M.-W.W. initiated the project, supervised the studies, analyzed the data, and wrote the manuscript with inputs from all co-authors; P.W. supervised the EM studies.

## References

1. Barberis C, Mouillac B, Durroux T. Structural bases of vasopressin/oxytocin receptor function. J Endocrinol 1998; 156:223–229.

2. Donaldson ZR, Young LJ. Oxytocin, vasopressin, and the neurogenetics of sociality. Science 2008; 322:900–904.

3. Tahara A, Saito M, Sugimoto T et al. Pharmacological characterization of the human vasopressin receptor subtypes stably expressed in Chinese hamster ovary cells. Br J Pharmacol 1998; 125:1463–1470.

4. Moeller HB, Rittig S, Fenton RA. Nephrogenic Diabetes Insipidus: Essential Insights into the Molecular Background and Potential Therapies for Treatment. Endocr Rev 2013; 34:278–301.

5. Keegan BP, Akerman BL, Pequeux C, North WG. Provasopressin expression by breast cancer cells: implications for growth and novel treatment strategies. Breast Cancer Res Treat 2006; 95:265–277.

6. North WG. Gene regulation of vasopressin and vasopressin receptors in cancer. Exp Physiol 2000; 85 Spec No:27S–40S.

7. Sinha S, Dwivedi N, Tao S et al. Targeting the vasopressin type-2 receptor for renal cell carcinoma therapy. Oncogene 2020; 39:1231–1245.

8. Zaoral M, Kolc J, Sorm F. Amino Acids and Peptides. LXXI. Synthesis of 1-Deamino-8-D-γ-Aminobutyrine Vasopressin, 1-Deamino-8-D-Lysine Vasopressin, and 1-Deamino-8-D-Arginine Vasopressin. Collect Czech Chem Commun 1967; 32:1250–1257.

9. Sebti Y, Rabbani M, Sadeghi HM, Sardari S, Ghahremani MH, Innamorati G. Effect of mutations in putative hormone binding sites on V2 vasopressin receptor function. Res Pharm Sci 2015; 10:259–267.

10. Slusarz MJ, Slusarz R, Ciarkowski J. Investigation of mechanism of desmopressin binding in vasopressin V2 receptor versus vasopressin V1a and oxytocin receptors: molecular dynamics simulation of the agonist-bound state in the membrane-aqueous system. Biopolymers 2006; 81:321–338.

11. Slusarz MJ, Gieldon A, Slusarz R, Ciarkowski J. Analysis of interactions responsible for vasopressin binding to human neurohypophyseal hormone receptors-molecular dynamics study of the activated receptor-vasopressin-G(alpha) systems. JPept Sci 2006; 12:180–189.

12. Waltenspuhl Y, Schoppe J, Ehrenmann J, Kummer L, Pluckthun A. Crystal structure of the human oxytocin receptor. Sci Adv 2020; 6:eabb5419.

13. Nehme R, Carpenter B, Singhal A et al. Mini-G proteins: Novel tools for studying GPCRs in their active conformation. Plos One 2017; 12.

14. Maeda S, Qu Q, Robertson MJ, Skiniotis G, Kobilka BK. Structures of the M1 and M2 muscarinic acetylcholine receptor/G-protein complexes. Science 2019; 364:552–557.

15. Kim K, Che T, Panova O et al. Structure of a Hallucinogen-Activated Gq-Coupled 5-HT2A Serotonin Receptor. Cell 2020; 182:1574–1588 e1519.

16. Ballesteros JA, Weinstein H. Integrated methods for the construction of three-dimensional models and computational probing of structure-function relations in G protein-coupled receptors. Methods in neurosciences 1995; 25:366–428.

17. Duan J, Shen DD, Zhou XE et al. Cryo-EM structure of an activated VIP1 receptor-G protein complex revealed by a NanoBiT tethering strategy. Nat Commun 2020; 11:4121.

18. Zhou F, Zhang H, Cong Z et al. Structural basis for activation of the growth hormone-releasing hormone receptor. Nat Commun 2020; 11:5205.

19. Du Vigneaud V, Ressler C, Trippett S. The sequence of amino acids in oxytocin, with a proposal for the structure of oxytocin. J Biol Chem 1953;205:949–957.

20. Lubecka EA, Sikorska E, Sobolewski D, Prahl A, Slaninova J, Ciarkowski J. Potent Antidiuretic Agonists, Deamino-Vasopressin and Desmopressin, and Their Inverso Analogs: NMR Structure and Interactions With Micellar and Liposomic Models of Cell Membrane. Biopolymers 2016; 106:245–259.

21. Chan WY, Wo NC, Stoev ST, Cheng LL, Manning M. Discovery and design of novel and selective vasopressin and oxytocin agonists and antagonists: the role of bioassays. Exp Physiol 2000; 85 Spec No:7S–18S.

22. Conner M, Hawtin SR, Simms J et al. Systematic analysis of the entire second extracellular loop of the V(1a) vasopressin receptor: key residues, conserved throughout a G-protein-coupled receptor family, identified. J Biol Chem 2007; 282:17405–17412.

23. Zhou Q, Yang D, Wu M et al. Common activation mechanism of class A GPCRs. Elife 2019; 8.

24. Wacker D, Wang C, Katritch V et al. Structural Features for Functional Selectivity at Serotonin Receptors. Science 2013; 340:615–619.

25. Pandy-Szekeres G, Munk C, Tsonkov TM et al. GPCRdb in 2018: adding GPCR structure models and ligands. Nucleic Acids Res 2018; 46:D440–D446.

26. Rasmussen SG, DeVree BT, Zou Y et al. Crystal structure of the beta2 adrenergic receptor-Gs protein complex. Nature 2011; 477:549–555.

27. Erdélyi L, Szalai L, Sziráki A, Balla A, Hunyady L. Functional characterization of inherited S127F substitution in V2 vasopressin receptor revealed a loss-of-function mutation leading to nephrogenic diabetes insipidus. Endocrine Abstracts 2017; 49.

28. Wenkert D, Schoneberg T, Merendino JJ, Jr. et al. Functional characterization of five V2 vasopressin receptor gene mutations. Mol Cell Endocrinol 1996; 124:43–50.

29. Chini B, Mouillac B, Balestre MN et al. Two aromatic residues regulate the response of the human oxytocin receptor to the partial agonist arginine vasopressin. FEBS Lett 1996; 397:201–206.

30. Wisniewski K, Galyean R, Tariga H et al. New, potent, selective, and short-acting peptidic V1a receptor agonists. J Med Chem 2011; 54:4388–4398.

31. Carpenter B, Nehme R, Warne T, Leslie AG, Tate CG. Structure of the adenosine A2A receptor bound to an engineered G protein. Nature 2016; 538:542.

32. Makita N, Manaka K, Sato J, Iiri T. V2 vasopressin receptor mutations. Vitam Horm 2020; 113:79–99.

33. Feldman BJ, Rosenthal SM, Vargas GA et al. Nephrogenic syndrome of inappropriate antidiuresis. N Engl JMed 2005; 352:1884–1890.

34. Carpentier E, Greenbaum LA, Rochdi D et al. Identification and characterization of an activating F229V substitution in the V2 vasopressin receptor in an infant with NSIAD. J Am Soc Nephrol 2012; 23:1635–1640.

35. Erdelyi LS, Mann WA, Morris-Rosendahl DJ et al. Mutation in the V2 vasopressin receptor gene, AVPR2, causes nephrogenic syndrome of inappropriate diuresis. Kidney Int 2015; 88:1070–1078.

36. Arthus MF, Lonergan M, Crumley MJ et al. Report of 33 novel AVPR2 mutations and analysis of 117 families with X-linked nephrogenic diabetes insipidus. J Am Soc Nephrol 2000; 11:1044–1054.

37. Bichet DG, Birnbaumer M, Lonergan M et al. Nature and recurrence of AVPR2 mutations in X-linked nephrogenic diabetes insipidus. Am J Hum Genet 1994; 55:278–286.

38. Fujiwara TM, Morgan K. Molecular-Biology of Diabetes-Insipidus. Annu Rev Med 1995; 46:331–343.

39. Schulz A, Grosse R, Schultz G, Gudermann T, Schoneberg T. Structural implication for receptor oligomerization from functional reconstitution studies of mutant V2 vasopressin receptors. J Biol Chem 2000;275:2381–2389.

40. Liu J, Wess J. Different single receptor domains determine the distinct G protein coupling profiles of members of the vasopressin receptor family. J Biol Chem 1996; 271:8772–8778.

41. Dehoorne JL, Raes AM, van Laecke E, Hoebeke P, Vande Walle JG. Desmopressin toxicity due to prolonged half-life in 18 patients with nocturnal enuresis. The Journal of urology 2006; 176:754–757; discussion 757-758.

42. Wisniewski K, Qi S, Kraus J et al. Discovery of Potent, Selective, and Short-Acting Peptidic V2 Receptor Agonists. JMed Chem 2019; 62:4991–5005.

43. Laskowski RA, Swindells MB. LigPlot+: multiple ligand-protein interaction diagrams for drug discovery. J Chem Inf Model 2011; 51:2778–2786.

44. Pettersen EF, Goddard TD, Huang CC et al. UCSF ChimeraX: Structure visualization for researchers, educators, and developers. Protein Sci 2020.

45. Koehl A, Hu H, Maeda S et al. Structure of the micro-opioid receptor-Gi protein complex. Nature 2018; 558:547–552.

46. Kang Y, Kuybeda O, de Waal PW et al. Cryo-EM structure of human rhodopsin bound to an inhibitory G protein. Nature 2018; 558:553–558.

47. Liang YL, Zhao P, Draper-Joyce C et al. Dominant Negative G Proteins Enhance Formation and Purification of Agonist-GPCR-G Protein Complexes for Structure Determination. ACS pharmacology& translational science 2018; 1:12–20.

48. Zhao LH, Ma S, Sutkeviciute I et al. Structure and dynamics of the active human parathyroid hormone receptor-1. Science (New York, NY) 2019; 364:148–153.

49. Dixon AS, Schwinn MK, Hall MP et al. NanoLuc Complementation Reporter Optimized for Accurate Measurement of Protein Interactions in Cells. ACS chemical biology 2016; 11:400–408.

50. Rasmussen SG, DeVree BT, Zou Y et al. Crystal structure of the β2 adrenergic receptor-Gs protein complex. Nature 2011; 477:549–555.

51. Mastronarde DN. Automated electron microscope tomography using robust prediction of specimen movements. Journal of structural biology 2005; 152:36–51.

52. Zivanov J, Nakane T, Scheres SHW. Estimation of high-order aberrations and anisotropic magnification from cryo-EM data sets in RELION-3.1. Iucrj 2020; 7:253–267.

53. Zheng SQ, Palovcak E, Armache JP, Verba KA, Cheng Y, Agard DA. MotionCor2: anisotropic correction of beam-induced motion for improved cryo-electron microscopy. Nat Methods 2017; 14:331–332.

54. Rohou A, Grigorieff N. CTFFIND4: Fast and accurate defocus estimation from electron micrographs. J Struct Biol 2015; 192:216–221.

55. Kucukelbir A, Sigworth FJ, Tagare HD. Quantifying the local resolution of cryo-EM density maps. Nat Methods 2014; 11:63–65.

56. Sanchez-Garcia R, J Gomez-Blanco, A Cuervo, JM Carazo, COS Sorzano, Vargas J. DeepEMhancer: a deep learning solution for cryo-EM volume post-processing. bioRxiv 2020.

57. Waterhouse A, Bertoni M, Bienert S et al. SWISS-MODEL: homology modelling of protein structures and complexes. Nucleic Acids Res 2018; 46:W296–W303.

58. Pettersen EF, Goddard TD, Huang CC et al. UCSF Chimera--a visualization system for exploratory research and analysis. J Comput Chem 2004; 25:1605–1612.

59. Emsley P, Cowtan K. Coot: model-building tools for molecular graphics. Acta Crystallogr D Biol Crystallogr 2004; 60:2126–2132.

60. Adams PD, Afonine PV, Bunkoczi G et al. PHENIX: a comprehensive Python-based system for macromolecular structure solution. Acta Crystallogr D Biol Crystallogr 2010; 66:213–221.

61. Croll TI. ISOLDE: a physically realistic environment for model building into low-resolution electrondensity maps. Acta Crystallogr D 2018; 74:519–530.

62. Sastry GM, Adzhigirey M, Day T, Annabhimoju R, Sherman W. Protein and ligand preparation: parameters, protocols, and influence on virtual screening enrichments. J Comput Aid Mol Des 2013; 27:221–234.

63. Genheden S, Ryde U. The MM/PBSA and MM/GBSA methods to estimate ligand-binding affinities. Expert Opin Drug Dis 2015; 10:449–461.

