## Supplementary Information for "Molecular basis of ligand recognition and activation of human V2 vasopressin receptor"

### Supplementary Figures

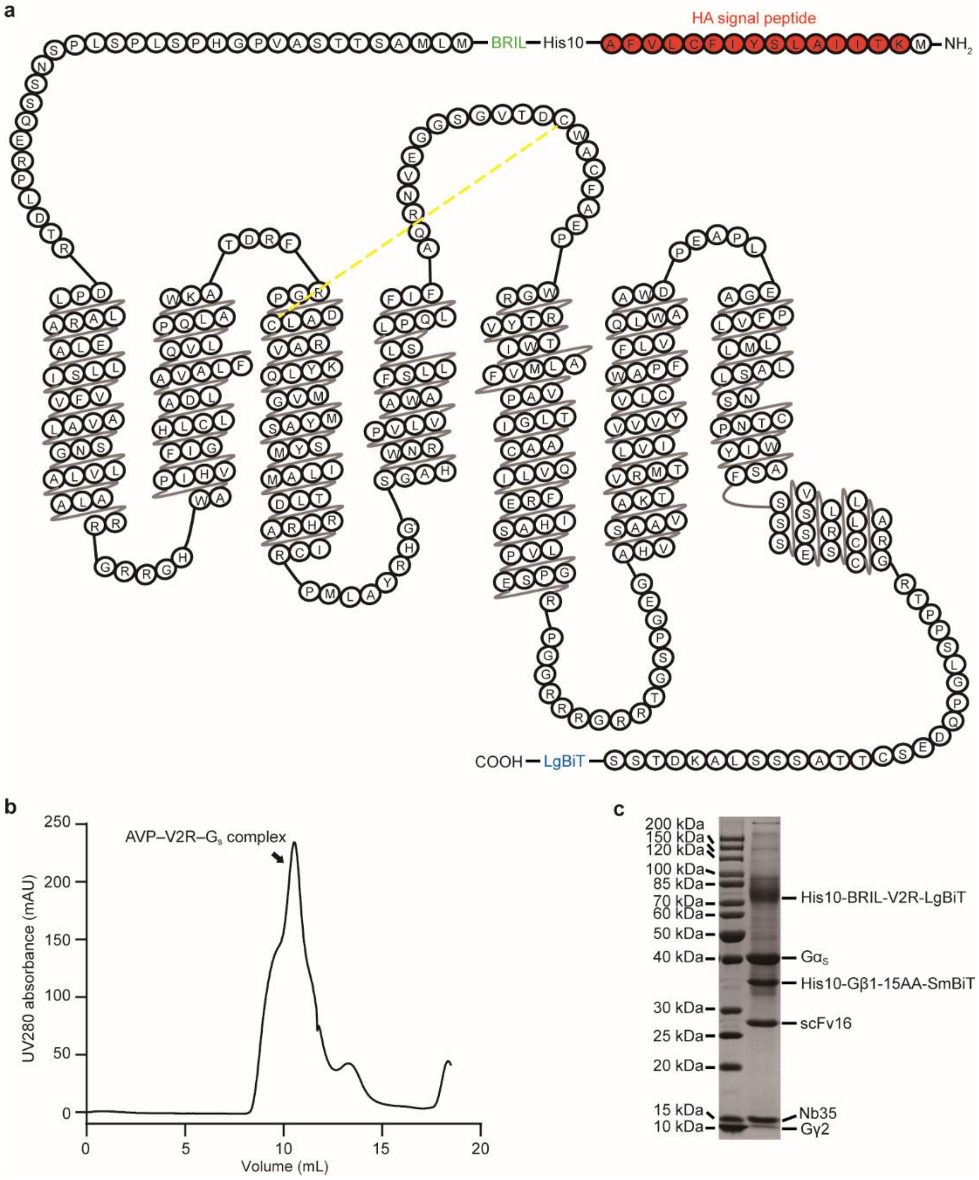


**Fig. S1.** Purification and characterization of the AVP–V2R–$G_S$ complex.

**a**, Schematic of HA–His10–BRIL–V2R(1-371)–LgBiT construct used in the study. The (HA) signal peptide (red), BRIL (green) and LgBiT (blue) are highlighted and indicated. The conserved disulfide bond between C112$^{3.25}$ and C192$^{ELC2}$ is depicted as yellow dashed lines. **b**, Representative size-exclusion chromatography elution profile of His-purified complex on Superdex 200 Increase 10/300 column. The monomer peak for AVP–V2R–$G_S$ complex is indicated. **c**, SDS-PAGE analysis of the purified complex concentrated from the monomeric fraction by Coomassie blue staining.

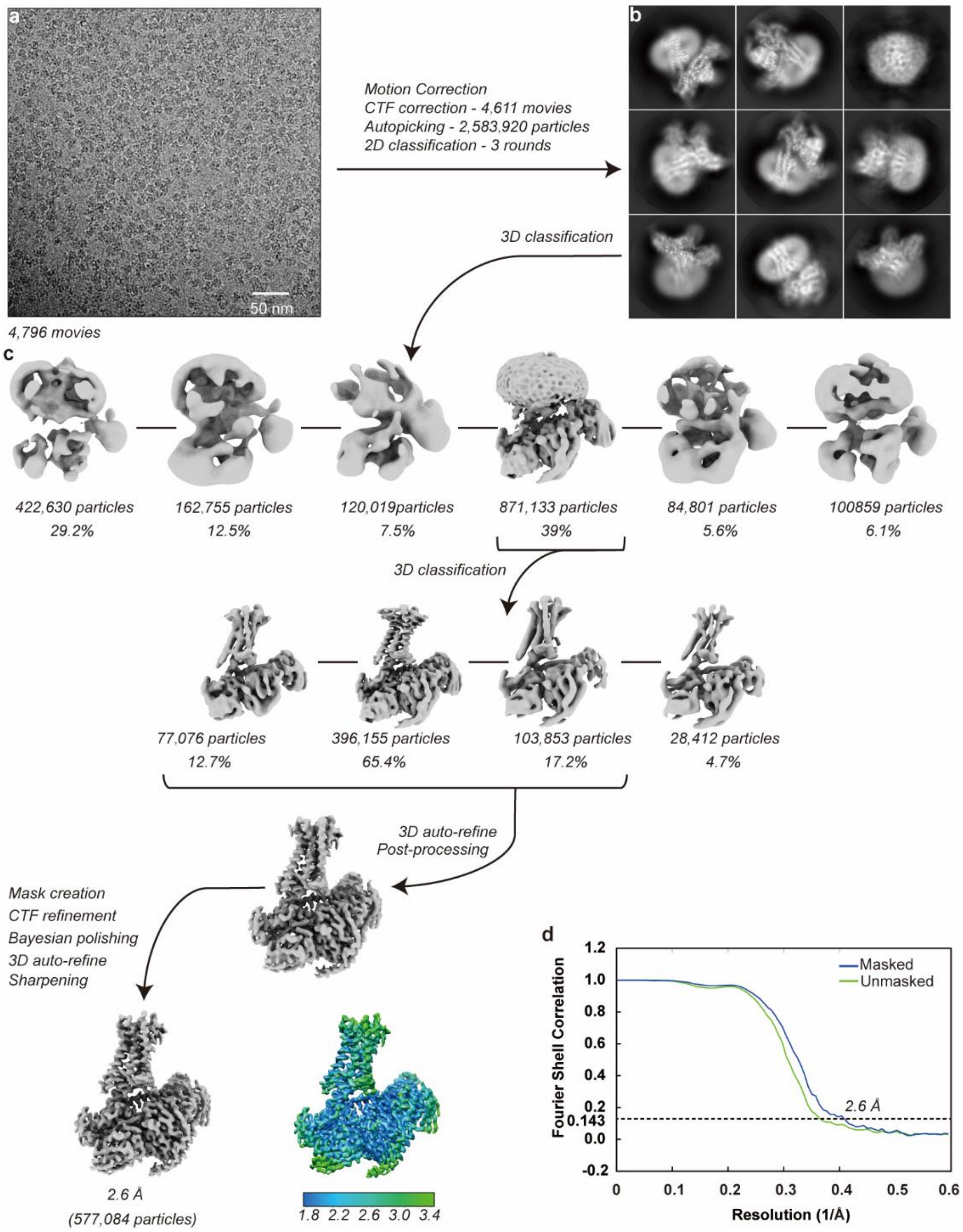


**Fig. S2.** Cryo-EM images and single-particle reconstruction of the AVP–V2R–$G_s$ complex. **a**, Representative cryo-EM micrograph of AVP–V2R–$G_s$ complex in the data collection. Scale bar, 50 nm. **b**, Represnetative 2D average classes showing distinguishable secondary structure features. **c**, Flow chart of cryo-EM data processing using Relion 3.1. **d**, The “Gold-standard” Fourier shell correlation curve indicates that the overall resolution of the electron density map of the complex is 2.6 Å.

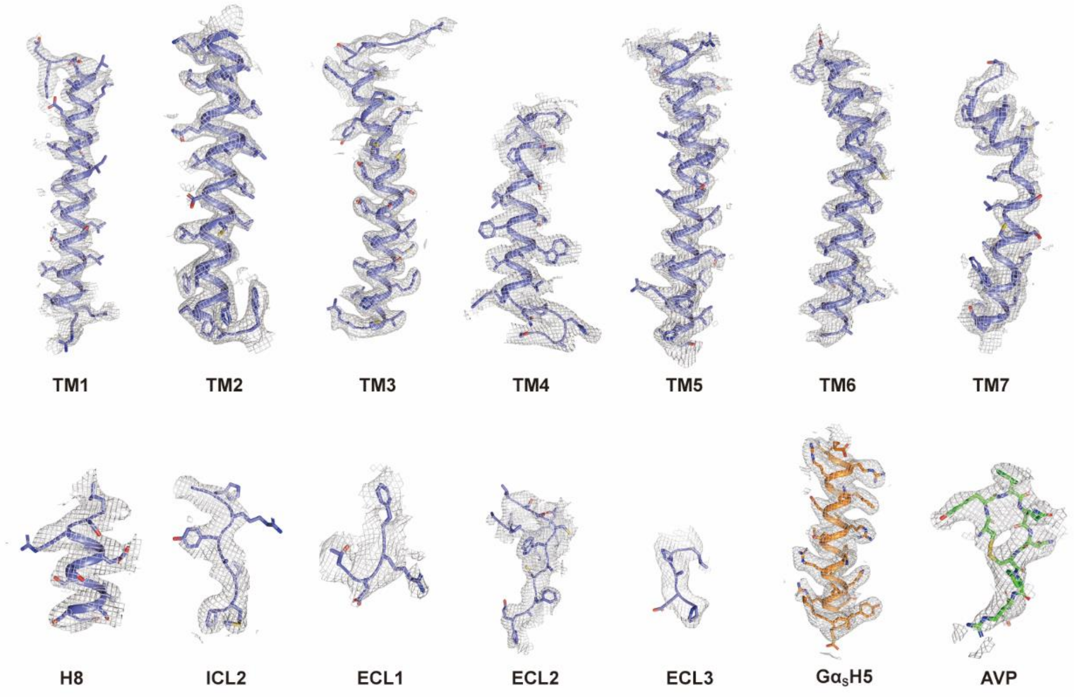


**Fig. S3.** Representative cryo-EM density map of the AVP–V2R–$G_s$ complex.

Cryo-EM density of the seven transmembrane (TM) helices, helix 8 (H8), AVP, ICL2, ECL1, ECL2, ECL3 of V2R, and α5 helix of $G\alpha_s$ ($G\alpha_s$H5). The densities of most side chains are clearly seen in the map.

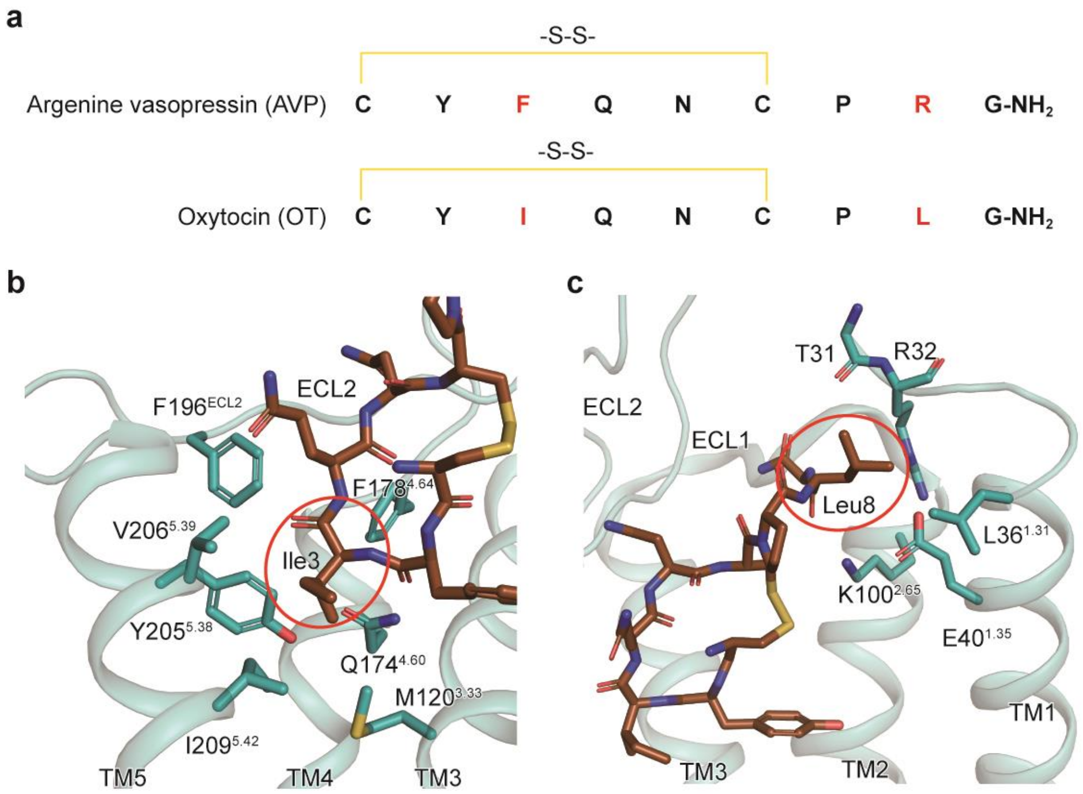


**Fig. S4.** Docking pose of OT in V2R.

**a,** Sequence alignment of argenine vasopressin (AVP) and oxytocin (OT). The third and eighth residues are different and colored in red. The disulfide bond between Cys1 and Cys6 are depicted as yellow lines. **b** and **c**, The predicted binding pose of OT in V2R. The receptor and the OT are shown as cartoon (teal) and sticks (brown), respectively. Residues Ile3 and Leu8 of OT are circled. Hydrogen bond is indicated as a red line.

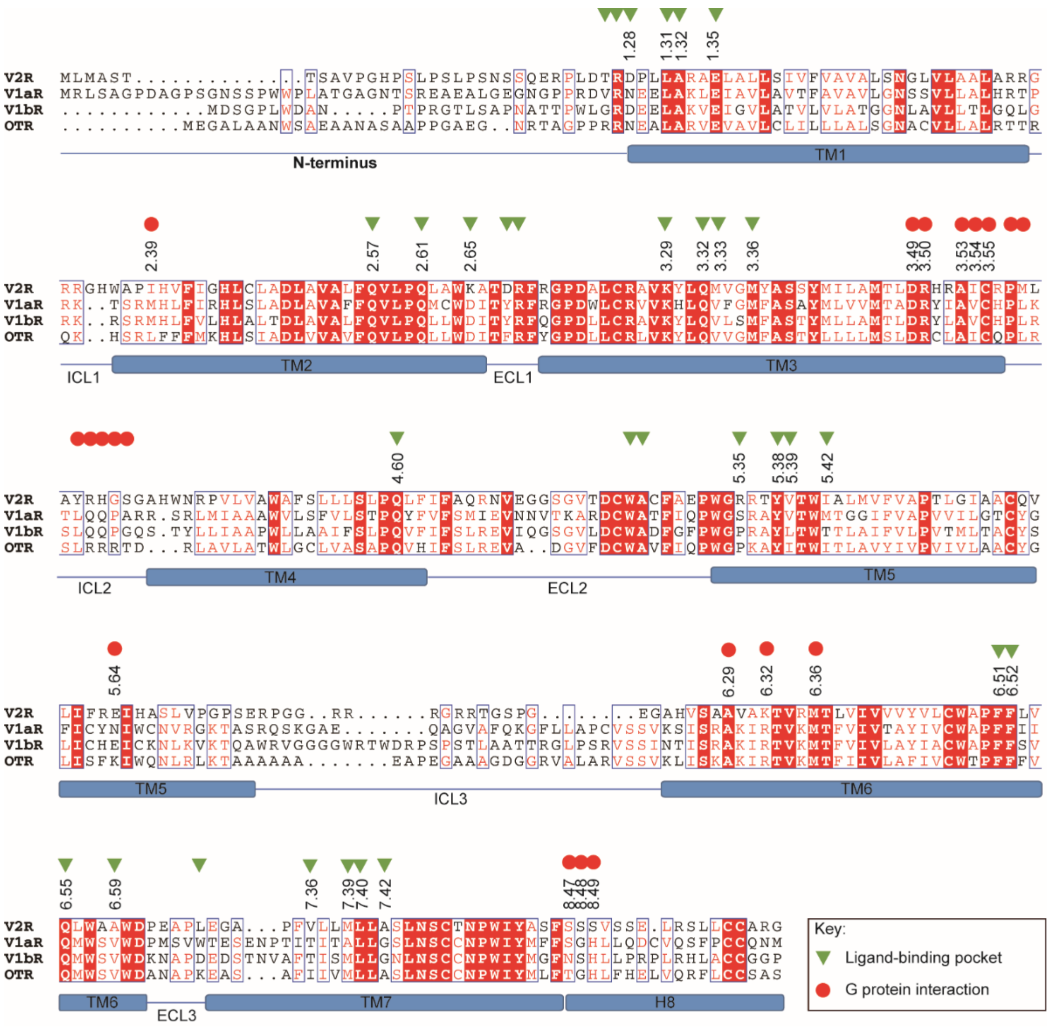


**Fig. S5.** Sequence alignment of the vasopressin/oxytocin receptor subfamily.

The overall sequence (except the C terminus) alignment of vasopressin/oxytocin receptor subfamily was generated using UniProt (http://www.uniprot.org/align/) and the graphics was prepared on the ESPript 3.0 server (http://espript.ibcp.fr/ESPript/cgi-bin/ESPript.cgi). Colors represent the similarity of residue: red background, identical; red text, strongly similar. Residues consisting of the ligand-binding pocket and $G_s$-coupling interface are indicated by green triangles and red circles, respectively. Secondary structure elements predicted by the GPCRdb (http://www.gpcrdb.org) are annotated underneath the sequences.

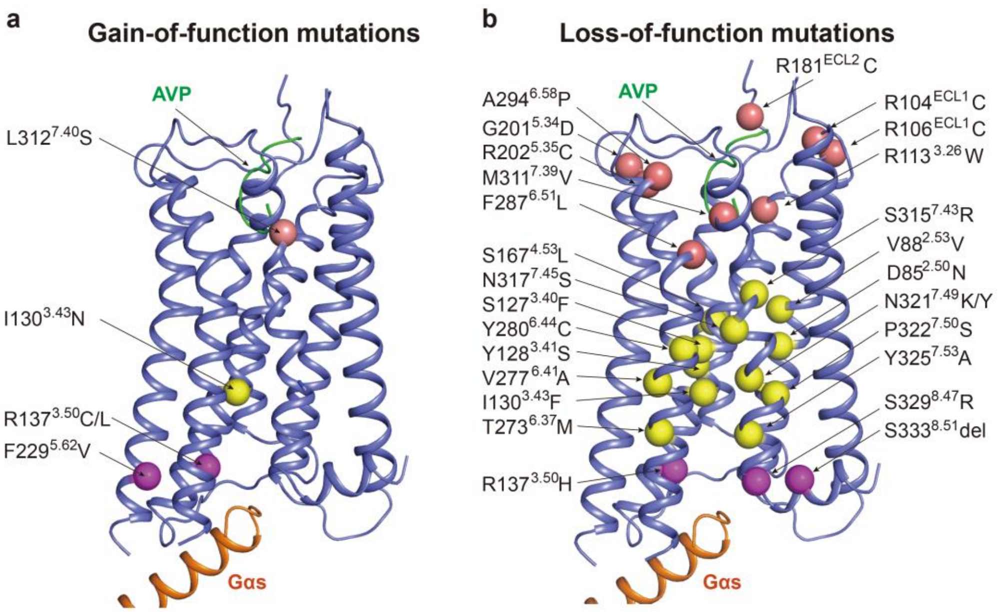


**Fig. S6.** The disease-associated V2R mutations.

Location of the gain-of-function (**a**) and loss-of function mutations (**b**) are indicated in the V2R structure (slate blue). Mutations situated in the ligand-binding pocket are shown as single salmon spheres, while those located in the G protein-coupling region are displayed as magenta spheres. Mutations in the central region connecting the ligand-binding pocket and G protein-coupling sites are indicated as yellow spheres. AVP (arginine vasopressin, green) and $G\alpha_s$ (orange) are presented as cartoon.

### Supplementary Tables

**Table S1.** Cryo-EM data collection, model refinement and validation statistics.

| | **AVP-V2R-Gs** |
|---|---|
| **Data collection and processing** | |
| Magnification | 165,000 |
| Voltage (kV) | 300 |
| Electron exposure ($e^-$/Å$^2$) | 60 |
| Defcous range (μm) | -0.5 ~ -2.0 |
| Pixel size (Å) | 0.42 |
| Symmetry imposed | C1 |
| Initial particle projections (no.) | 2,583,920 |
| Final particle projections (no.) | 577,084 |
| Map resolution (Å) | 2.6 |
| FSC threshold | 0.143 |
| Map resolution range (Å) | 1.9-3.4 |
| **Refinement** | |
| Initial model used | 6TPK |
| Model resolution (Å) | 2.8 |
| FSC threshold | 0.5 |
| Map sharpening *B* factor (Å) | -59.70 |
| Model composition | |
| Non-hydrogen atoms | 10,102 |
| Protein residues | 1,283 |
| Ligand | 1 |
| Lipids | 3 |
| *B* factors (Å$^2$) | |
| Protein | 111.89 |
| Lipids | 134.06 |
| RMSD | |
| Bond lengths (Å) | 0.012 |
| Bond angles (°) | 1.037 |
| Validation | |
| MolProbity score | 0.90 |
| Clashscore | 1.00 |
| Rotamer outliers (%) | 0.09 |
| Ramachandran plot | |
| Favored (%) | 97.46 |
| Allowed (%) | 2.54 |
| Disallowed (%) | 0 |

RMSD, root-mean-square deviation.

**Table S2.** Effects of construct modifications on AVP-induced cAMP accumulation.

| Construct | cAMP | | Cell-surface expression (% WT) |
|---|---|---|---|
| | $pEC_{50}$ ± S.E.M. | $E_{max}$ (%WT) | |
| V2R(1-371) (WT) | 11.95 ± 0.03 | 100 | 100 |
| His10-BRIL-V2R(1-371)-LgBiT | 12.03 ± 0.05 | 99.51 ± 0.20 | 51.47 ± 11.65** |

cAMP accumulation and cell-surface expression assays were performed in HEK 293T cells expressing wild-type (WT) V2R (1-371) or His10-BRIL-V2R(1-371)-LgBiT (construct for cryo-EM study), respectively. cAMP data were analyzed using a three-parameter logistic equation to determine $pEC_{50}$ and $E_{max}$. The $E_{max}$ and expression values were normalized to the WT which was set to 100%. All data are shown as means ± S.E.M. of at least three independent experiments. One-way ANOVA was used to determine the *P* value compared with the response of the WT. **P*<0.01; ***P*<0.001; ****P*<0.0001.

**Table S3.** Effects of alanine scanning mutagenesis of the residues in the ligand-binding pocket on AVP-induced cAMP accumulation.

| Mutants | cAMP | | Cell-surface expression (% WT) |
|---|---|---|---|
| | $pEC_{50}$ ± S.E.M. | $E_{max}$ (% WT) | |
| Wild-type | 11.73±0.07 | 100 | 100 |
| T31A | 11.69±0.06 | 99.09±0.83 | 79.95±9.64 |
| R32A | 11.32±0.11 | 98.11±1.38 | 86.46±9.47 |
| D33A | 11.57±0.05 | 99.61±0.83 | 106.02±5.53 |
| L36A | 10.56±0.05*** | 100.11±1.65 | 83.77±6.73 |
| E40A | 11.12±0.10** | 98.20±0.71 | 53.08±5.93** |
| Q92A | 10.51±0.14*** | 98.20±2.55 | 89.47±2.33 |
| Q96A | 9.85±0.09*** | 99.05±2.13 | 95.32±12.71 |
| K100A | 10.81±0.02*** | 98.99±1.86 | 89.84±8.08 |
| D103A | 11.63±0.09 | 97.94±2.16 | 125.76±3.14 |
| Q119A | 10.43±0.04*** | 100.16±1.42 | 84.54±2.80 |
| M120A | 11.25±0.09 | 99.27±1.17 | 75.77±7.46 |
| M123A | 11.17±0.16 | 99.38±1.46 | 65.06±4.01 |
| Q174A | 9.31±0.15*** | 101.26±0.85 | 55.21±4.02** |
| D191A | 10.38±0.22*** | 99.63±0.84 | 88.49±10.06 |
| W193A | 11.26±0.29 | 99.65±1.63 | 108.73±5.58 |
| R202A | 11.73±0.15 | 98.11±0.75 | 85.73±11.94 |
| Y205A | nd | nd | 34.87±3.13 |
| V206A | 11.24±0.13 | 99.69±1.72 | 74.38±6.68 |
| I209A | 11.39±0.01 | 100.37±1.39 | 69.90±8.87 |
| F288A | 10.50±0.09*** | 99.50±0.06 | 15.56±0.71*** |
| Q291A | 11.94±0.06 | 99.40±0.99 | 69.04±4.72 |
| L302A | 11.69±0.06 | 100.80±0.78 | 75.76±4.59 |
| V308A | nd | nd | 0.32±0.06*** |
| M311A | 11.55±0.02 | 100.37±0.09 | 55.63±7.23** |
| L312A | 11.39±0.07 | 99.15±0.51 | 50.60±13.50** |

nd, not determined.

$pEC_{50}$ and $E_{max}$ values were determined using a three-parameter logistic equation. All data shown are as means ± S.E.M. from at least three independent experiments. $E_{max}$ and cell-surface expression values were normalized to the wild-type (WT) which was set to 100%. One-way ANOVA was used to determine the *P* value compared with the response of the WT. *$P<0.01$; **$P<0.001$; ***$P<0.0001$.

**Table S4.** Interaction of AVP with residues in the V2R ligand-binding pocket.

| Vasopressin | V2R | Interaction |
|---|---|---|
| Cys1 | Q96$^{2.61}$ K116$^{3.29}$ | Backbone-side chain hydrogen bond |
| | M311$^{7.39}$ | Hydrophobic interaction |
| Tyr2 | Q92$^{2.57}$ Q96$^{2.61}$ Q119$^{3.32}$ | Hydrophobic interaction |
| | Q174$^{4.60}$ | Backbone-side chain hydrogen bond |
| | F287$^{6.51}$ M311$^{7.39}$ | Hydrophobic interaction |
| | L312$^{7.40}$ | Side chain-backbone hydrogen bond |
| | A314$^{7.42}$ | Hydrophobic interaction |
| Phe3 | M120$^{3.33}$ M123$^{3.36}$ Y205$^{5.38}$ V206$^{5.39}$ I209$^{5.42}$ F287$^{6.51}$ F288$^{6.52}$ Q291$^{6.55}$ | Hydrophobic interaction |
| Gln4 | R202$^{5.35}$ Q291$^{6.55}$ | Side chain-side chain hydrogen bond |
| | Q291$^{6.55}$ | Side chain-backbone hydrogen bond |
| | A295$^{6.59}$ | Hydrophobic interaction |
| Asn5 | W193$^{ECL2}$ | Hydrophobic interaction |
| | A194$^{ECL2}$ | Side chain-backbone hydrogen bond |
| Cys6 | K100$^{2.65}$ | Hydrophobic interaction |
| Pro7 | L302$^{ECL3}$ V308$^{7.36}$ | Hydrophobic interaction |
| Arg8 | R32 | Hydrophobic interaction |
| | D33$^{1.28}$ | Side chain-side chain hydrogen bond |
| | L36$^{1.31}$ A37$^{1.32}$ | Hydrophobic interaction |
| | E40$^{1.32}$ | Electrostatic interaction |
| Gly9 | T31 R32 D103$^{ECL1}$ R104$^{ECL1}$ | Hydrophobic interaction |
| $NH_2$ | D103$^{ECL1}$ | Side chain-backbone hydrogen bond |
| | R104$^{ECL1}$ | Hydrophobic interaction |
